# Pho92 direct interaction with Not1 recruits the CCR4-NOT complex to m^6^A containing mRNAs to modulate meiosis in *S. cerevisiae*

**DOI:** 10.64898/2026.10.27.762167

**Authors:** Lina Sène, Ditipriya Hazra, Jérémy Scutenaire, Nelly Morellet, Nathalie Ulryck, Mélody Matelot, Halil Bounoua, Mouna Raji, Sana Afensiss, Bastien Morlet, Carine van Heijenoort, Bertrand Séraphin, Marc Graille

**Author notes:** Correspondence should be addressed to BS or MG. These authors contributed equally (Lina Sène, Ditipriya Hazra). RNA Molecular Biology Group, National Institute of Arthritis and Musculoskeletal and Skin Disease, NIH, Bethesda, MD 20892, USA. Department of Biotechnology, St. Xavier’s College, Kolkata, 700016, India.

## Abstract

N^6^-methyladenosine (m^6^A) is the most abundant internal modification in eukaryotic mRNAs. In *S. cerevisiae*, m^6^A is exclusively deposited during meiosis, where it plays an essential role in meiotic progression. In this context, the absence of Pho92, the sole known m^6^A-binding protein in budding yeast, leads to the abnormal persistence of m^6^A-containing mRNAs, ultimately resulting in delayed meiosis. Here, we investigate how m^6^A recognition by Pho92 regulates gene expression. Using complementary *in vivo* and *in vitro* approaches, we identify direct interactions between Pho92 and Not1, the scaffold subunit of the CCR4-NOT deadenylase complex. Structural analyses reveal that a short evolutionarily conserved motif in Pho92 adopts an α-helical conformation that docks onto specific regions of Not1. Functionally, these interactions are required for efficient meiotic progression. Altogether, our findings demonstrate that Pho92 couples m^6^A recognition to the main deadenylase to shape the meiotic transcriptome. More broadly, our work provides mechanistic insights into how epitranscriptomic signals interface with the core mRNA decay machinery to regulate gene expression during cellular differentiation.

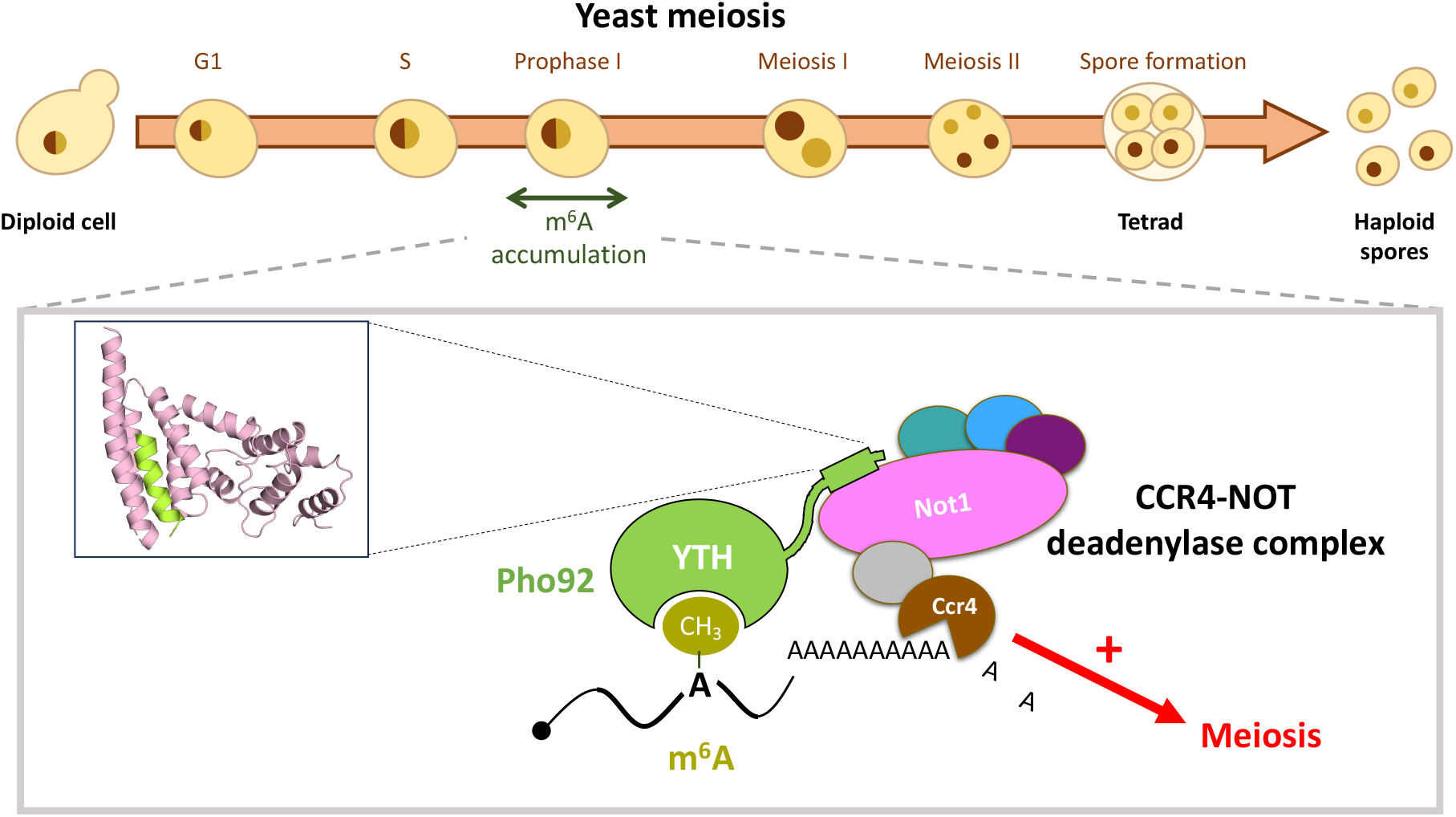

## Introduction

The life cycle of eukaryotic messenger RNAs (mRNAs) is a tightly orchestrated process, encompassing synthesis, maturation, cytoplasmic export, translation, and degradation. Each of these steps is subject to intricate regulatory controls that allow cells to fine-tune protein production in response to internal and external cues. Such regulation is particularly critical during cell division (both mitosis and meiosis) for which precise coordination of gene expression ensures faithful genome transmission during cell duplication or gamete formation. These mechanisms often involve synergistic actions, such as the combined modulation of transcription and mRNA decay rates, to rapidly induce or repress the expression of specific proteins (1).

mRNA turnover is a central mechanism in gene expression control. The stability of individual transcripts is influenced by a myriad of factors that monitor their structural and sequence features, thereby setting basal and regulated decay rates. In particular, the CCR4-NOT complex integrates various signals present in individual mRNAs to modulate the kinetics of their poly(A) tail shortening and subsequent elimination (2). Over the past decade, the epitranscriptomic mark N6-methyladenosine (m^6^A) has emerged as a new pivotal player in regulating mRNA decay. In human cells, m^6^A modifications are co-transcriptionally deposited on nascent mRNAs by a large holoenzyme complex, primarily composed of the catalytic subunit METTL3 and its partner METTL14, along with several co-factors (3–5). In the yeast *Saccharomyces cerevisiae*, the orthologous m^6^A writer complex, termed the MIS complex, includes the catalytic subunit Ime4 (ortholog of METTL3) proposed to be associated with non-catalytic subunits including Mum2, Kar4 (ortholog of METTL14), Vir1, Slz1, and Dyn2 (6–9).

In metazoa, the presence of m^6^A on mRNAs has been reported to profoundly impact nearly every stage of the mRNA life cycle, from splicing and nuclear export to translation and stability (5, 10), emphasizing the pleiotropic role of m^6^A in gene expression regulation. Modified residues are recognized specifically by m^6^A reader proteins, among which the YTH domain-containing proteins are the most well-characterized (11–13). YTH proteins typically function by recruiting effector molecules that modulate the fate of m^6^A-modified transcripts or by sterically blocking access to specific mRNA sites. Notably, the recognition of m^6^A by cytoplasmic YTH proteins has been repeatedly linked to mRNA decay (14–18). Given the central role of m^6^A and its readers on the control of gene expression, they have been shown to influence a broad spectrum of physiological processes, including embryogenesis, circadian rhythms, stem cell differentiation, brain development, and carcinogenesis (19, 20).

One of the cellular processes widely influenced by m^6^A is meiosis. Mutations disrupting m^6^A deposition cause gametogenesis defects in both mice (21) and fruit flies (22), highlighting its conserved role. In *S. cerevisiae*, this role is also critical: several components of the m^6^A writer complex, including Ime4, are expressed during early meiosis and are essential for meiotic onset (7–9, 23, 24). Following meiotic induction, m^6^A levels on mRNAs peak at approximately 5 hours and then decline (6, 7, 25–28), a dynamic pattern consistent with active regulation. Importantly, not only is the deposition of m^6^A crucial for meiosis, but so is its recognition by Pho92 (also known as Mrb1 for methylated RNA-binding protein 1), the sole YTH protein in yeast (27, 28). Pho92 is transiently expressed during early meiosis and is pivotal for ensuring timely meiotic progression (26–28). Through direct binding to m^6^A marks, Pho92 appears to accelerate the downregulation of its mRNA targets (27, 28).

Pho92 was originally suggested to regulate genes involved in the yeast phosphate signal transduction (PHO) pathway in a process involving mRNA decay and the CCR4-NOT complex (29). However, these initial observations are puzzling given the specific expression of Pho92 during meiosis, when m^6^A is exclusively present (6, 7, 25–28). If the implication of Pho92 in meiosis is solidly established, the precise molecular mechanisms by which Pho92 modulates meiotic gene expression remain incompletely understood. This is due in part to the dynamic and profound remodeling of the transcriptome during meiosis, which poses significant challenges for assessing the stability of individual transcripts. Here, we have identified the molecular basis for Pho92’s interaction with the CCR4-NOT complex. Through biochemical and structural analyses, we pinpoint a short evolutionarily conserved motif in Pho92 that is critical for its binding to a central region of Not1, the scaffolding protein of the CCR4-NOT complex. Structural analyses and modeling detail how this motif folds into an α helix that docks directly on Not1. We further demonstrate that this interaction and is required for efficient meiosis. Altogether, our data show that Pho92 controls the meiotic gene expression program by recruiting the CCR4-NOT deadenylase. This work not only sheds light on the epitranscriptomic control of meiosis but also provides broader insights into post-transcriptional regulation of gene expression during cellular differentiation.

## Materials and Methods

### Yeast strains

All yeast strains used in this study are listed in Supplementary **Table S1** and are derived from the sporulation-proficient SK1 background with CUP1-inducible promoter that drives HA-tagged IME1 expression (pCUP-3HA-IME1; (30)). Strains for two-hybrid assays were derived from MAV203 (Invitrogen). The lithium acetate transformation protocol was used to introduce exogenous plasmids, oligonucleotides (listed in **Supplementary Tables S2 and S3**, respectively) or PCR-amplified linear DNA fragments (31).

Deletions and point mutations were generated using the CRISPR/Cas9 system (32). The absence of the deleted fragment or the presence of the desired mutation was confirmed by PCR. For point mutations, their presence was confirmed by the introduction of silent polymorphic restriction site. The sequence of the mutated region was determined by sequencing PCR products.

Pho92 was N-terminally tagged with TAP using the tagging cassette from pBS1761 ((33), **Supplementary Table S2**). Not1 was C-terminally tagged with His (3HA-HisMX) using the tagging cassette from pFA6a-3HA-HisMX6 ((34), **Supplementary Table S2**).

Diploids were obtained by transforming haploid cells with a YCp50 plasmid carrying the HO endonuclease gene and a URA3 marker (gift from Ed Hurt). The presence of the *MAT*a and *MATα* genes was verified by PCR on isolated colonies. The HO-containing plasmid was lost in rich medium before further use of the strains.

To generate specific genetic combinations, haploid yeast strains were crossed on YPDA medium to generate heterozygous diploids, which were selected using genomic or plasmid markers. After plasmid loss, diploids were sporulated for 2 days at 30 °C on media containing 50 µM CuSO₄. Asci were digested with Zymolyase in SCE buffer (1 M sorbitol, 0.1 M sodium citrate, pH 7) for 15 min. 10 µL of suspension were plated on YPDA (1% yeast extract, 2% peptone, 2% glucose supplemented with 10 mg/L adenine hemisulfate salt), and tetrads were dissected with a Singer MSM system. Isolated spores were grown for 2 days at 30 °C on YPDA, and their genotypes were determined by replica plating on selective media. Mating type was determined by crosses with reference strains (BSY1174 *MATα* and BSY1177 *MAT*a), and allele segregation was confirmed by PCR when required.

### Growth conditions and meiosis induction

Cells in vegetative growth were grown in YPDA at 30 °C with shaking at 170 rpm. Synchronous meiotic entry was performed as follows: respiratory-proficient diploids were preselected on 1% yeast extract, 2% peptone, 3% glycerol, 2% agar plates for 24 h at 30 °C, transferred for an additional 24 h of growth in YPD (1% yeast extract, 2% peptone, 4% dextrose) at 30 °C with shaking at 170 rpm, diluted in BYTA medium (1% yeast extract, 2% tryptone, 1% potassium acetate, 50 mM potassium phthalate) to OD_600_ = 0.2 and incubated for an additional 16 h at 30 °C and 170 rpm. Next, cells were washed once with sterile water and resuspended in SPO medium (0.3% potassium acetate) at OD_600_ = 2.0 and incubated at 30 °C and 170 rpm. Because we used the pCUP-3HA-IME1 background, 50 µM of copper (II) sulfate was added after 2 h in SPO.

### Meiotic progression with DAPI counting

300 µL of sporulating cells (OD_600_ = 2.0) were collected at the indicated time points, centrifuged for 2 min at 8,000 g, fixed in 1 volume of 70% ethanol, and stored at 4 °C until analysis. Cells were then pelleted and resuspended in 2 µg/mL DAPI solution in water for 5 min before analysis. Meiotic divisions were monitored by counting the number of DAPI masses per cell in 200 cells per time point, with a Leica DM 4000B microscope.

### Sequence alignment

To align Pho92 sequences, the Pho92 protein sequences from a subset of *Saccharomycetaceae* species were retrieved from the NCBI database using keyword and BLAST sequence searches. Pho92 sequences were aligned with Clustal Omega (35), and alignments were manually edited. Colored alignments were exported using Jalview (36).

### Immunoprecipitation and Mass Spectrometry

PMSF (final concentration of 1 mM) was added to 100 mL of meiotic cultures incubated in sporulation media for 5 hours. After 2 min of PMSF treatment at room temperature with head-over-tail rotation, cells were collected by centrifugation for 2 min at 5,000 g and 4 °C. Cells were washed once with sterile water containing 1 mM PMSF, collected by centrifugation as above and the pellet was snap-frozen in liquid nitrogen. Frozen pellets were resuspended in 300 µL of lysis buffer (150 mM KCl, 10 mM Tris pH 8.0, 0.1% IGEPAL (Sigma), 1 mM PMSF, 1× Complete mini protease inhibitors (Roche), 2 µM Pepstatin A) and homogenized with glass beads at 4 °C once for 40 sec at 6 m/s using a MP Bio FastPrep-24 Cell Homogenizer. Cell debris were pelleted by centrifugation for 5 min at 5,000 g and 4 °C. The supernatant was transferred to a new tube and centrifuged for 5 min at 15,000 g and 4 °C. Total protein concentration was determined using the Bradford assay (Biorad). 300 µL of supernatant was mixed with 50 µL of IgG paramagnetic beads (Dynabeads^TM^ Thermofischer scientific) previously equilibrated with lysis buffer. Samples were incubated with head-over-tail rotation for 30 min at 4 °C. Beads were washed three times with 1 mL of lysis buffer. For western Blot analysis, samples were eluted with Laemmli buffer (60 mM Tris-HCl pH 6.8, 10% glycerol, 0.002% bromophenol blue, 2% sodium dodecyl sulfate and 5% dithiothreitol) by boiling the samples. For mass spectrometry analyses, three additional washes were performed with 1% PBS and the dried beads were snap-frozen before trypsin digestion. Biological triplicates were performed for each condition.

Mass spectrometry analyses were performed by the IGBMC Proteomic platform. Samples were injected in technical triplicates on the QExactive instrument. All raw files were analyzed in MaxQuant v1.6.0.13 against the 2019 SwissProt *S. cerevisiae* protein database & additional data analyses were performed in Perseus v1.4.0.2.

### Western blotting and antibodies

1 mL of meiotic cells or YPDA-grown cells (OD_600_ = 2.0) was collected by centrifugation for 2 min at 8,000 g and room temperature and the pellet was stored at -80 °C until use. Total proteins were extracted as described previously (37). Proteins samples were separated by SDS-PAGE and transferred onto Amersham™ Protran™ 0.45 µm nitrocellulose membranes (GE Healthcare, Cat# 10600033) at 4 °C for 1-2 h and 110 V. After Ponceau staining, membranes were blocked in 5% milk in PBS-T (1× PBS buffer with 0.1% Tween-20) and incubated for 1 h at room temperature with one of the following antibodies: anti-HA (1:2,000; Eurogentec, Cat# MMS-101P-500), anti-Stm1 (1:10,000; kind gift from F. Wyers). After four washes with PBS-T, membranes were incubated for 1 h at room temperature with Peroxidase-AffiniPure Goat Anti-Mouse (1:5,000; Jackson ImmunoResearch, Cat# 115-035-003) or Anti-rabbit IgG, HRP-linked antibody (1:5,000; Cell Signaling, Cat# 7074S). TAP-tagged proteins were detected after a single incubation of the membranes for 1 h at room temperature with Peroxidase Anti-Peroxidase Complex produced in rabbit (1:2,000; Sigma, Cat# P1291). Chemiluminescence signals were detected using an ImageQuant 800 (Cytiva) apparatus with Immobilon® Crescendo Western HRP substrate (Millipore, Cat# WBLUR0500) according to the manufacturer’s instructions.

### Yeast 2-hybrid assay

The ProQuest two-hybrid system was used to assay protein-protein interactions. The desired inserts were amplified by PCR, transferred into the pDONR221 plasmid backbone using the Gateway cloning strategy (Invitrogen) and verified by sequencing. Point mutations were introduced by site-directed mutagenesis into the Gateway Entry plasmid pDONR221 derivative and validated by sequencing. Inserts were then transferred into the relevant backbones (pDEST22 and/or pDEST32, **Supplementary Tables S2**). ß-galactosidase activities were determined using the Beta-Glo Assay system (Promega) according to the manufacturer’s recommendation. Luminescence signals were measured using a Centro XS LB 960 luminometer (Berthold). The average values and standard error of the mean (SEM) of five biological replicates were plotted.

### Cloning, expression and purification of recombinant proteins

The gene fragments encoding for the domains of interest from Pho92, Not1 or Caf40 proteins were amplified using *S. cerevisiae* genomic DNA by PCR and inserted into entry plasmids listed in **Supplementary Table S4**. To improve the expression levels of Not1 domains, we expressed them fused to a N-terminal tag consisting of an His_6_ tag followed by two copies of the Z domain from the IgG binding *Staphylococcus aureus* protein A (His_6_-ZZ tag; (38)). First, we inserted a *de novo* synthesized DNA sequence (gbMG12; Integrated DNA Technologies, Belgium) encoding for the His_6_-ZZ tag followed by a 3C protease cleavage site into the pET28b plasmid using *Nco*I and *BamH*I restriction enzymes to generate the pET28b-His_6_-ZZ-3C plasmid. Next, the PCR products encompassing various Not1 domains were cloned into the latter backbone using *EcoR*I and *Not*I restriction enzymes (Thermo Fisher Scientific; **Supplementary Table S4**). Plasmids encoding for the Pho92 or Not1 mutants were obtained by one-step site-directed mutagenesis using oligonucleotides and plasmids listed in **Supplementary Table S4**.

The pMG965 and pMG966 plasmids allowing the expression of ScNot1(1343–1565)-(GSS)_5_-ScPho92(21–44) and ScPho92(21–44)-(GSS)_5_-ScNot1(1343–1565), respectively, were obtained by inserting *de novo* synthesized DNA fragments (gbMG21 and gbMG22, respectively; Integrated DNA Technologies, Belgium) into the pET28b-His_6_-ZZ-3C plasmid using *BamH*I and *Xho*I restriction enzymes (See **Supplementary Table S4**). Information regarding the oligonucleotides used for amplification, the entry plasmids, the restriction enzymes used, the names of resulting plasmids and the fusion protein expressed are listed in **Supplementary Table S4**.

Wild-type (WT) and mutant derivatives of the GST-Pho92(1–69) fusion proteins were expressed in *E. coli* BL21 (DE3) Codon+ cells after transformation with appropriate plasmid (**Supplementary Table S4**). Cultures were performed in 1 L of terrific broth auto-inducible media (ForMedium AIMTB0260) containing ampicillin (100 µg/mL) and chloramphenicol (25 µg/mL) first for 3 h at 37 °C and then overnight at 18 °C. The cells were harvested by centrifugation at 3,300 g for 30 min. The pellet was resuspended in 30 mL lysis buffer L1 (20 mM Tris-HCl pH 7.5, 200 mM NaCl, 5 mM β-mercaptoethanol, 1 mM MgCl_2_) in the presence of 500 µM PMSF. Cell lysis was performed by sonication, followed by lysate clearance by centrifugation at 20,000 g for 30 min. The supernatants were applied on GSH-sepharose resin pre-equilibrated with lysis buffer L1. After extensive washing steps with lysis buffer L1 supplemented with 0.01 mM ATP, the protein was eluted with elution buffer (20 mM Tris-HCl pH 7.5, 200 mM NaCl, 10 mM GSH, 5 mM β-mercaptoethanol). The eluted protein was next subjected to HiTrap Q FF ion-exchange chromatography and eluted with a linear gradient of NaCl from 50 mM to 1 M NaCl (20 mM Tris-HCl pH 7.5, 5 mM β-mercaptoethanol). Proteins of interest were finally injected onto a S75 16/60 size-exclusion chromatography (SEC) column pre-equilibrated with SEC buffer (20 mM Tris-HCl pH 8, 150 mM NaCl, 5 mM β-mercaptoethanol).

The different Not1 constructs were expressed using the same conditions used for the GST-Pho92 fragments with the exception that the ampicillin antibiotics was replaced by kanamycin (100 µg/mL). The bacteria were resuspended in lysis buffer L2 (20 mM Tris-HCl pH 8, 200 mM NaCl, 5 mM β-mercaptoethanol) in the presence of 500 µM PMSF. After cell lysis and clarification as described above, Not1 constructs were purified on Protino NiNTA beads equilibrated with lysis buffer L2. After elution with elution buffer (20 mM Tris-HCl pH 8, 200 mM NaCl, 5 mM β-mercaptoethanol, 400 mM imidazole pH 7), the proteins were loaded onto an HiTrap Q FF ion-exchange chromatography and eluted with a linear gradient of NaCl from 50 mM to 1 M NaCl (20 mM Tris-HCl pH 8, 5 mM β-mercaptoethanol). Proteins of interest were finally injected onto a S75 16/60 size-exclusion chromatography column pre-equilibrated with SEC buffer. When the removal of the His_6_-ZZ tag was necessary, the proteins eluted from the NiNTA resin were incubated overnight with human rhinovirus 3C protease under dialysis condition against dialysis buffer (20 mM Tris-HCl pH 8, 100 mM KCl, 5 mM β-mercaptoethanol).

The Not1-(GSS)_5_-Pho92 and Pho92-(GSS)_5_-Not1 fusion proteins were expressed as described above for GST-Pho92 using pMG965 and pMG966, respectively. The proteins were purified as described for GST-Pho92(1–69) proteins with minor differences. Specifically, after elution from GSH-sepharose resin, the GST tag was removed by overnight incubation with human rhinovirus 3C protease under dialysis condition against buffer (20 mM Tris-HCl pH 8, 100 mM NaCl, 5 mM β-mercaptoethanol). The proteins were then loaded on HiTrap Heparin column and eluted as described above. Finally, the dialysis buffer was used to equilibrate the S75-16/60 column. The SeMet-labeled Pho92-(GSS)_5_-Not1 fusion protein was expressed using auto-inducible SeMet labeling medium supplemented with ampicillin at 100 μg/mL and chloramphenicol at 25 μg/mL as described (39) and purified as the native protein.

To express the His_6_-SUMO-Caf40(102–373)/His_6_-ZZ-Not1(1071–1282) complex, the pMG1247 plasmid encoding for His_6_-SUMO-Caf40(102–373) was transformed in *E. coli*ArcticExpress (DE3) cells. These cells were grown in 1L of LB media containing kanamycin (100 µg/mL) and gentamicin (15 µg/mL), first for 3 h at 30 °C and then protein expression was induced by addition of 0.5 mM IPTG and transfer of the bacteria at 13 °C for 18 h. After centrifugation at 3,300 g for 30 min, the bacteria were resuspended in lysis buffer L3 (50 mM Tris-HCl pH 8, 300 mM NaCl, 5 mM β-mercaptoethanol). One bacterial pellet was mixed with one bacterial pellet of *E. coli* cells expressing the His_6_-ZZ-Not1(1071–1282) protein (obtained as indicated above). Prior to cell lysis by sonication, 500 µM PMSF, 5 mM MgCl_2_, 1 mM DTT and lysozyme (30 mg) were added to the lysis buffer. After cell lysis and clarification as described above, proteins of interests were loaded on Protino NiNTA beads equilibrated with lysis buffer L3. The beads were washed first with 50 mM Tris pH 8, 300 mM NaCl, 5 mM β-mercaptoethanol, 10 mM MgCl_2_, 10 mM ATP and then 50 mM Tris pH 8, 300 mM NaCl, 5 mM β-mercaptoethanol, 20 mM imidazole pH 7. The complex was then eluted from the Protino NiNTA beads using 50 mM Tris pH 8, 300 mM NaCl, 5 mM β-mercaptoethanol, 400 mM imidazole pH 7 and diluted into a buffer containing 50 mM Tris pH 8, 5 mM β-mercaptoethanol to reach a final NaCl concentration of 50 mM prior injection on HiTrap Q FF ion-exchange chromatography column. The proteins were eluted with a linear gradient of NaCl from 50 mM to 1 M NaCl (50 mM Tris-HCl pH 8, 5 mM β-mercaptoethanol). Proteins of interest were finally injected onto a S200 16/60 size-exclusion chromatography column pre-equilibrated with SEC buffer (50 mM Tris-HCl pH 8, 3000 mM NaCl, 5 mM β-mercaptoethanol).

### Crystallization, data processing and structure determination

Diffracting crystals were obtained by mixing equal volumes of purified Pho92-(GSS)_5_-Not1 fusion protein at 37 mg/mL in the dialysis buffer and crystallization condition (1.2M Na/K phosphate pH 7, 1mM Proline) at 4°C. Crystals were cryo-protected by transfer into the crystallization condition supplemented with 15% and then 30% ethylene glycol and then flash-frozen in liquid nitrogen.

Diffraction data were collected on beamline Proxima 2A at synchrotron SOLEIL (Saint-Aubin, France) and were processed with the XDS and AIMLESS programs (40, 41). Three datasets (space group C222(1)) were collected at inflection, peak and remote wavelengths of Se-edge from a single SeMet-labeled protein crystal to solve the structure by SeMAD (Multiple Anomalous Dispersion; see **Supplementary Table S5** for data collection statistics). Twenty Se sites (from four Pho92-(GSS)_5_-Not1 molecules) were successfully located using SHELXD (42) and experimental phasing was performed with SHARP (43). NCS-averaging and density modification were performed using the RESOLVE program (44). The resulting experimental map was used to manually build an initial model using COOT (45). This initial model was used as search model to locate 8 copies in a high-resolution dataset (space group C2, processed with STARANISO server (46)). The model was iteratively improved by cycling building and refinement steps performed with COOT and BUSTER (47), respectively (see **Supplementary Table S5** for refinement statistics). Structure coordinates have been deposited in the PDB database.

### GST pulldown

Pull-down experiments were performed by mixing 25 µg of GST-Pho92(1–69) proteins (WT or L32R/L35R), or GST alone as control, with 50 µg of the His_6_-ZZ-Not1(1071–1282) or His_6_-ZZ-Not1(1071–1282)/His_6_-SUMO-Caf40(102–373) complex. All purified proteins were free of nucleic acids according to the OD_280nm_/OD_260nm_ ratio. Binding buffer (20 mM Tris-HCl pH 7.5, 200 mM NaCl, 5 mM β-mercaptoethanol) was added to a final volume of 60 µL. The reaction mixtures were incubated on ice for 1 hour. 10 µL was withdrawn as « input » fraction. The remaining 50 µL were incubated at 4°C for 1-1.5 hours on a rotating wheel with 40 µL of glutathione paramagnetic agarose beads (Thermo) pre-equilibrated with binding buffer in a final volume of 200 µL. Beads were washed three times with 500 µL of binding buffer. Bound proteins were eluted with 50 µL of elution buffer (binding buffer supplemented with 20 mM reduced glutathione). 4 µL of « input » and 20 µL of « elution » fractions were resolved on SDS-PAGE and visualized by Coomassie blue staining. The presence of His_6_-ZZ-Not1(1071–1282) or His_6_-SUMO-Caf40(102–373) proteins was checked by western blot using anti-His antibodies coupled to HRP (1:4000; Sigma-Aldrich, Cat# #A7058).

### NMR spectroscopy

For ^15^N-^13^C labeling of the Not1(1343–1565) domain, the transformed bacteria were grown in 1L of M9 medium containing ^15^NH_4_Cl (1g/L final concentration) and ^13^C_6_ D-Glucose (4g/L final concentration). Protein expression was induced by addition of IPTG when the bacteria were transferred from 37 °C to 18 °C. The protein was purified as described for the unlabeled protein.

NMR experiments were performed at 293 K and 308 K on 800 MHz and 950 MHz AVANCE III HD Bruker spectrometers equipped with TCI cryoprobes. The backbone (N, H, CO, Cα, Cβ, Hα, Hβ) assignments were obtained using standard 3D triple resonance assignment experiments on a ^15^N-^13^C labeled protein sample: HN(CO)CACB, HNCACB, HNCA, HN(CO)CA, HNCO and HN(CA)CO to assign all the carbon atoms in a given residue using the CcpNmr Analysis 2.4.2 software (48). Backbone and side-chain atoms assignments were obtained using standard triple resonance assignment experiments on a ^15^N-^13^C labeled protein sample. Not1(1343–1565) assignments were deposited into BMRB. The mapping of the Pho92 binding site was performed by comparing the HSQC spectra of the ^15^N-^13^C labeled protein alone or in the presence of 1.5 molar excess of the peptide corresponding to Pho92 residues 21 to 44, at either 293 K or 308 K.

### ITC

ITC experiments were performed at 20 °C using an ITC-200 microcalorimeter (MicroCal). All proteins were prepared in 20 mM Tris-HCl pH 8, 150 mM NaCl, 5 mM β-mercaptoethanol. For all titration experiments, 20 injections of 2 µL of titrant (Not1 constructs at 1-1.1 mM) were added to the target (GST-Pho92(1–69) WT or mutants at 90-100 µM) at intervals of 180 s. The heat of dilution of the titrant was determined from the peaks measured after full saturation of the target by the titrant. The experimental data were fitted using the ORIGIN^®^ software assuming a one-binding site model.

### Fluorescence anisotropy measurements

The N-terminally FITC-labeled peptide (PM194) encompassing residues 21 to 44 from *S. cerevisiae* Pho92 was synthesized by the IGBMC Peptide Synthesis facility using the Fmoc chemistry. Crude peptide products were purified by reverse phase high performance liquid chromatography (HPLC). Quality and purity of the final products were assessed by HPLC and mass spectrometry.

Affinities between the PM194 peptide and the different Not1 constructs were quantified by fluorescence anisotropy using a FP8300 spectrofluorimeter (Jasco) at 20°C using excitation and emission wavelengths of 495 nm and 520 nm, respectively. Increasing amounts of Not1 constructs (0 to 30-50 µM) were added into a solution composed of 20 mM Tris-HCl pH 8, 100 mM NaCl, 5 mM 2-mercaptoethanol and containing the PM194 peptide at a concentration of 12.8 nM in a 1 mL final volume. Experiments were performed in triplicates. Kd and dR_max_ (maximum value of anisotropy difference) were calculated by fitting experimental curves with equation (A) using the program OriginLabs:

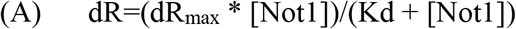

where dR is the anisotropy difference for a given Not1 concentration ([Not1]).

Competition experiments were performed by incubating Not1 domains (30 µM) with PM194 fluorescent peptide (12.8 nM) in the above conditions and then chasing the peptide by increasing concentrations of the GST-Pho92(1–69) (WT or L32R/L35R mutant). Dissociation constant (Kd_GST-Pho92_) was determined by fitting experimental curve with equation (B):

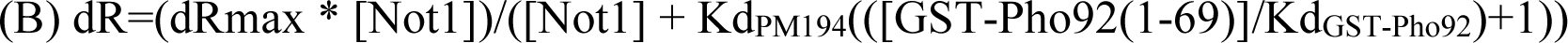

where dR is the anisotropy difference for a given concentration of GST-Pho92(1–69).

## Results

### The Pho92 N-terminal contains a conserved motif required for efficient sporulation

Our previous analyses demonstrated that Pho92 binds m^6^A residues deposited in mRNAs during sporulation and its deletion delays the meiotic program (27). Pho92 contains a canonical YTH domain at its C-terminus (residues 145-306) that was previously shown to bind m^6^A. Contrary to the YTH domain, which is conserved among eukaryotic cytoplasmic m^6^A binding proteins, the N-terminal region that precedes this domain, is predicted to be an intrinsically disordered region (IDR) and is not conserved between yeast and human. Because short sequence motifs evolve rapidly, we looked for conserved residues by aligning the full-length sequences of Pho92 homologs from different *Saccharomycetaceae* species. This revealed the presence of a conserved motif located between residues 21 and 44, in addition to the YTH domain (**Fig. 1A**). In the Pho92 IDR preceding the YTH domain, other residues were essentially not conserved (**Fig. S1**).

**Figure 1.**
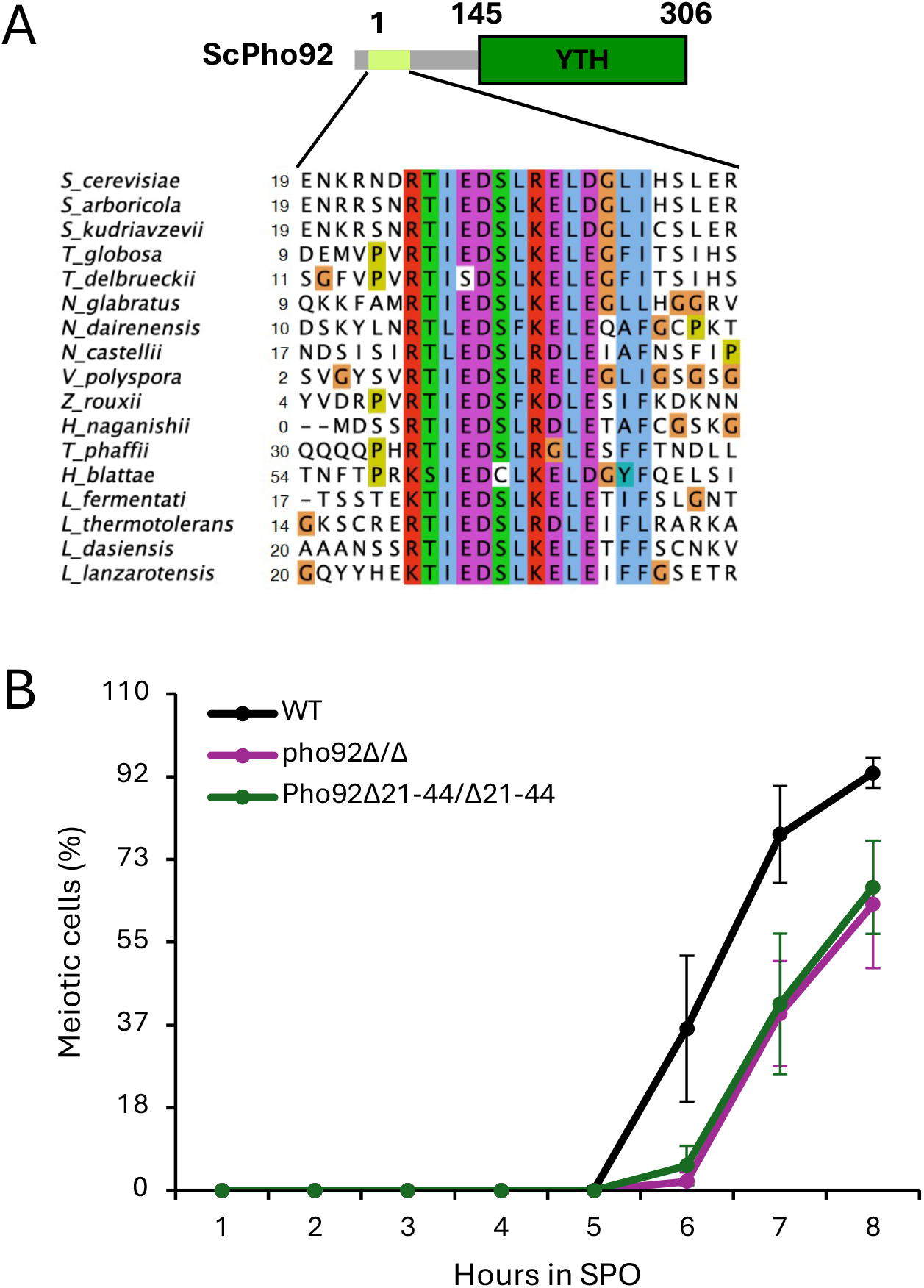
Pho92 contains a conserved N-terminal motif important for meiosis. **A.** Schematic representation of the *S. cerevisiae* Pho92 protein with the intrinsically disordered N-terminal region (residues 1-144) represented as a thick grey line and the C-terminal YTH domain (residues 145-306) colored in dark green rectangle. A conserved motif present in the Pho92 N-terminal region (**Suppl.** Fig. 1) is indicated as a light green rectangle and a multiple sequence alignment of this motif is presented below with coloring of conserved residues according to the Clustal Omega software (66). **B.** Comparison of the sporulation kinetics of homozygous *pho92Δ/Δ* and *pho92Δ21-44/Δ21-44* diploid strains versus the wild-type control (n=3). The absence of the conserved motif in Pho92 phenocopies the absence of Pho92.

To test whether this motif contributes to Pho92 function, we constructed an allele lacking this motif (*pho92(Δ21-44)*) and compared the sporulation kinetics of a homozygous diploid mutant with those of both a *pho92Δ/Δ* and a wild-type isogenic strain (**Fig. 1B**). As previously observed, deletion of PHO92 resulted in a meiotic delay compared to the wild-type strain. Strikingly, upon transfer to a sporulation medium, the *pho92Δ21-44/Δ21-44* strain exhibited the same extended meiosis kinetics as the *pho92Δ/Δ* strain (**Fig. 1B**). This observation indicates that this motif is functionally important for sporulation.

### Pho92 interacts with the CCR4-NOT complex

To identify partners of the Pho92 N-terminal motif, we prepared total protein extracts from a TAP-Pho92 strain 5 hours after meiosis induction, a time point corresponding to the peak of m^6^A modification in mRNAs, and immunoprecipitated the tagged protein using IgG beads. A wild-type strain without a tag served as a control for non-specific precipitation. To exclude factors that might bind Pho92 through its YTH domain, we also analyzed proteins co-purifying with the YTH domain alone using a TAP-Pho92(ΔN) strain. Proteins co-precipitating with the TAP-tagged factors were identified by mass spectrometry. Gene ontology (GO) term analysis revealed that the identified proteins were significantly (p<0.01) associated with components of cytoplasmic stress granules, (cytoplasmic) ribonucleoprotein granules, supramolecular complexes, CCR4-NOT complex, and membrane less organelles. Consistently, we identified different subunits of the CCR4-NOT complex namely Not1 (also named Cdc39), Not4 (Mot2), and Caf40, that were specifically enriched with TAP-Pho92 but under-represented in the untagged control (**Fig. 2A**). Strikingly, subunits of the CCR4-NOT complex were also underrepresented among proteins recovered from the TAP-Pho92(ΔN) construct (**Fig. 2A**), indicating that the Pho92 N-terminal region is required for this interaction.

**Figure 2.**
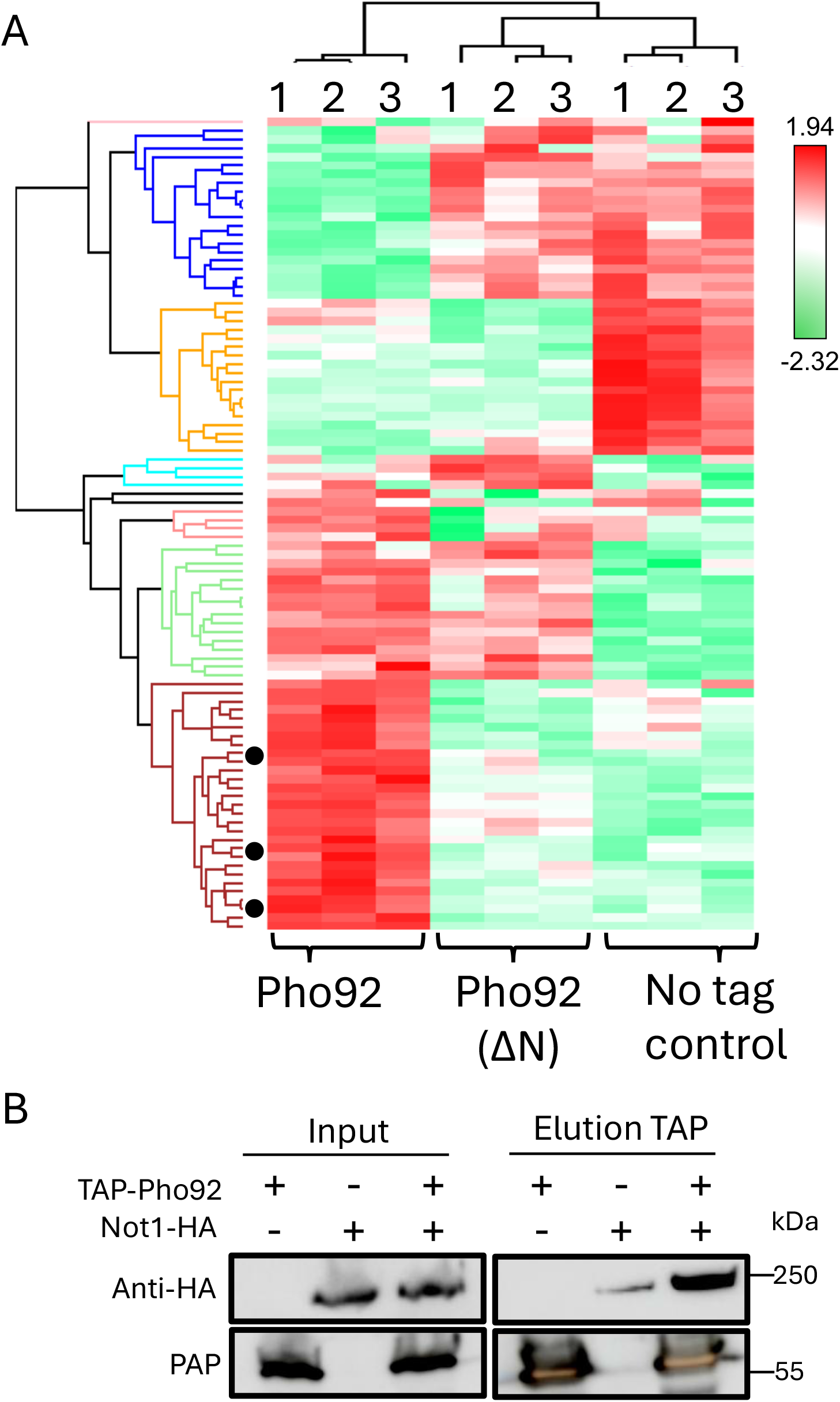
Pho92 interacts with the CCR4-NOT complex in a manner dependent on its N-terminal region. **A.** Heatmap of the differential levels of protein co-purified with TAP-Pho92, TAP-Pho92(ΔN) and from an untagged wild-type control strain as negative control. Proteins were identified by mass spectrometry from triplicate biological samples and an ANOVA (with Benjamini-Hochberg FDR=0.01) test was performed to pinpoint proteins significantly enriched in specific samples. 94 proteins were identified as significant, and from those proteins, a classic Euclidean Hierarchical clustering was made with protein clusters depicted in different colors on the left. The cluster in brown contains partners of Pho92 N-terminal region. GO term analysis revealed a significant presence of CCR4-NOT complex subunits (black circles): Caf40, Not1 (Cdc39) and Not4 (Mot2). **B.** Co-immunoprecipitation assay confirming the interaction between the Not1 and Pho92 proteins. Total protein extracts (input) were incubated with IgG Dynabeads and proteins co-immunoprecipitated with TAP-Pho92 were resolved by SDS-PAGE. Not1-HA and TAP-Pho92 were detected using anti-HA antibodies and the Peroxidase-Anti-Peroxidase (PAP) complex respectively.

To validate the interaction between Pho92 and the CCR4-NOT complex, we performed co-immunoprecipitation using a yeast strain carrying TAP- and HA-tagged chromosomal versions of Pho92 and Not1, respectively. We observed the specific co-immunoprecipitation of Not1 with Pho92 in extracts from meiotic *S. cerevisiae* cells (**Fig. 2B**), thereby confirming the interaction of Pho92 with the major deadenylase complex.

### Delineating Pho92 and Not1 regions required for their interaction

To more precisely characterize the regions of Not1 and Pho92 involved in their interaction, we performed two-hybrid assays. We first validated that Pho92 interacts with Not1 in this system. The Caf1 deadenylase (also known as Pop2), which was previously shown to interact directly with Not1 (49), served as a positive control. In these experiments, both Caf1 and Pho92 showed interaction with full-length Not1 (**Fig. 3A**). Using constructs encoding various Not1 regions, we detected an interaction between Pho92 and the C-terminal half of Not1 (Not1(1080–2108)), as well as a more defined region spanning residues 1071-1565 (**Fig. 3A**). No significant interaction was detected for other Not1 regions tested including the Not1(754–999) region, which interacted with Caf1 as expected. However, it is noteworthy that several Not1-derived constructs generated high and variable background values in this assay, potentially masking weak signals.

**Figure 3.**
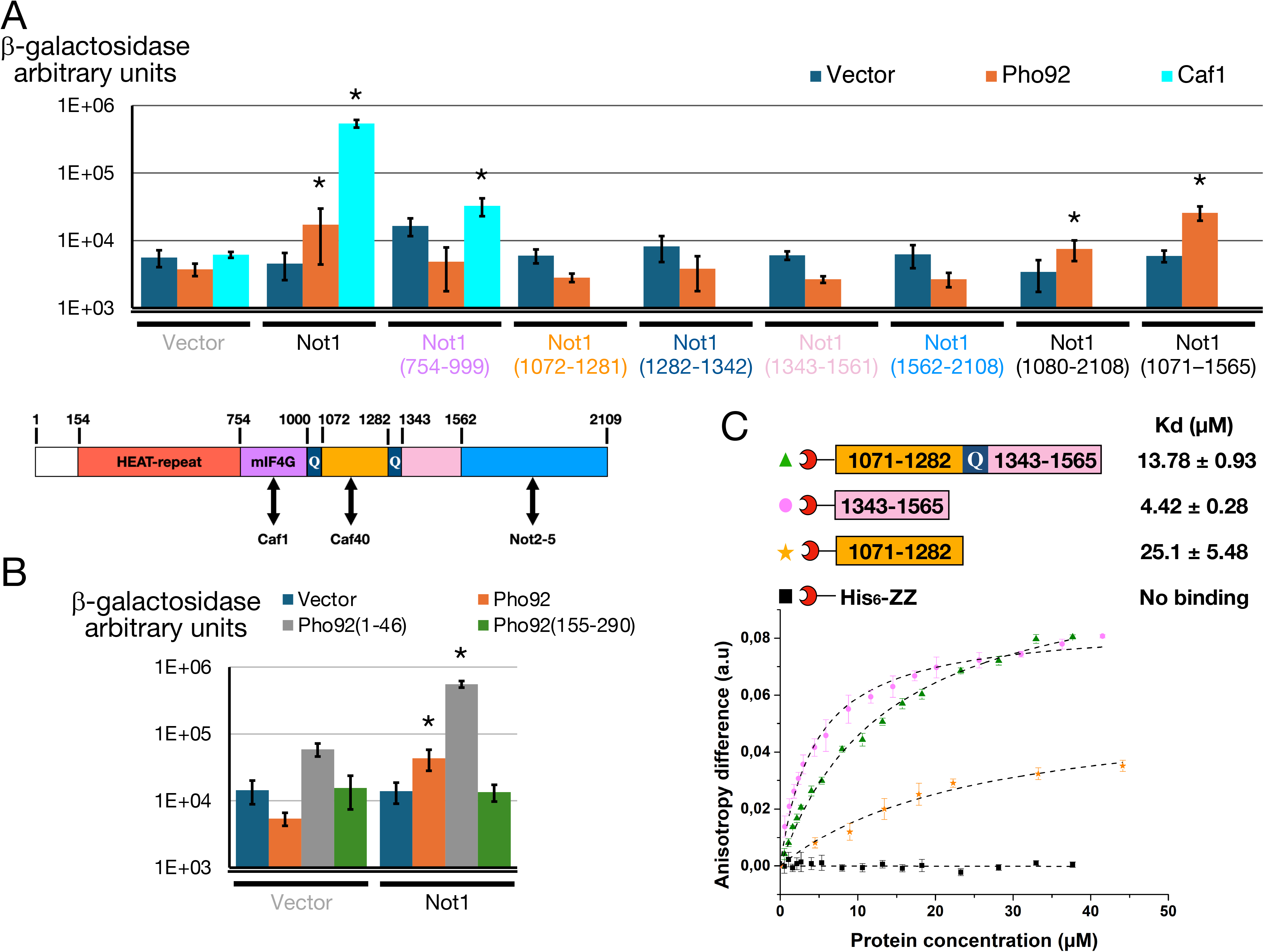
Pho92 possesses a conserved region needed for the interaction with domains of Not1. **A.** Interaction between Pho92 and Not1 or Not1 domains were assayed using the yeast two-hybrid assay. The Pho92 and Not1 coding sequences were fused to the DNA Binding Domain (DBD) or transcription Activation Domain (AD) respectively. Interactions were scored by assaying the β-galactosidase activity from 5 biological replicates. When average β-galactosidase activity value of the tested interaction was higher that the matching controls, a Kruskal-Wallis test validated non-homogeneity of the samples with a p-value < 0.05 (*). Schematic representation of the *S. cerevisiae* Not1 protein with the delineation of functional domains. The domain color code is used to color corresponding domains in the X axis of the β-galactosidase activity histogram. Known partners of functional domains are indicated by double arrows below the diagram. **B.** The N-ter region of Pho92 is necessary and sufficient to interact with Not1. Interaction between Pho92 domains and complete Not1 were monitored by yeast two-hybrid assay. Pho92 and Not1 coding sequences were fused to the DBD and AD domains respectively. Assays were performed as for panel A. **C.** Fluorescence anisotropy analyses indicate that Pho92(21–44) interacts with two domains in the Not1(1071–1565) region. The FITC-labeled Pho92(21–44) peptide was incubated with increasing amount of purified recombinant yeast His_6_-ZZ-Not1 domains. The graph represents the difference in anisotropy between the free fluorescent peptides and the reaction mix as a function of Not1 domains concentration. The curves obtained after fitting of the experimental data as described in Materials and Methods are shown as dashed lines. Error bars were calculated from triplicate experiments. The His_6_-ZZ tag used to purify the protein was used as a negative control. The Kd values determined from the fitting of the experimental data are indicated together with the schematic representation of the Not1 domains used.

In parallel, we prepared two constructs, each comprising one of the conserved regions of Pho92: the YTH domain (Pho92(155–290)) and the N-terminal motif (Pho92(1–46)). Consistent with the TAP-purification results (**Fig. 2A**), the YTH domain did not yield a positive signal with full-length Not1 in two-hybrid assays (**Fig. 3B**). In contrast, the Pho92 N-terminal motif alone was sufficient to mediate binding to Not1 (**Fig. 3B**).

Overall, the two-hybrid assays indicated that the region containing the short-conserved motif at the N-terminus of Pho92 interacts with the 1071-1565 region of Not1. Since these assays were performed in yeast, we cannot exclude the possibility that other endogenous factors bridged the interaction between Pho92 and Not1. However, since the smallest Not1 region sufficient to interact with Pho92 (residues 1071-1565) is known to bind only Caf40, this suggests that the interaction is most likely direct.

To more precisely define the regions involved in this interaction and confirm that a direct interaction between Pho92 and Not1, we quantified the binding between Pho92 residues 21-44 and several recombinant Not1 fragments within the region identified by two-hybrid assays. Using a fluorescent peptide corresponding to the conserved Pho92 region, we performed fluorescence anisotropy measurements. These revealed that the Not1(1071–1565) region interacts with this Pho92 peptide with a Kd value of 13.78 µM (**Fig. 3C**), constant with our two-hybrid assay data. We then purified two Not1 subdomains: Not1(1071–1282) and Not1(1343–1565). Surprisingly, both interacted with the Pho92(21–44) peptide (**Fig. 3C**). The Not1(1071–1282) domain (also known as CC domain), previously shown to interact with Caf40 (CNOT9 in human (50, 51)), binds the Pho92(21–44) peptide with a Kd of 25.1 µM. The poorly characterized Not1(1343–1565) domain, located downstream, interacts with higher affinity (Kd of 4.42 µM). All these interactions were specific, as we detected no binding to the fluorescent peptide of the His_6_-ZZ tag used for protein expression and purification (**Fig. 3C**).

Thus, using a combination of experimental approaches, we demonstrate that the conserved motif in the *S. cerevisiae* Pho92 m^6^A reader directly interacts with both the CC domain and the poorly characterized 1343-1565 region of Not1, the scaffolding protein of the CCR4-NOT complex.

### The N-terminal conserved motif of Pho92 adopts a helical structure and binds to a hydrophobic region on Not1(1343–1565) domain

To understand the molecular bases underlying the recognition of Pho92 by Not1(1343–1565), whose function and binding partners, to our knowledge, remain unidentified, we tried to reconstitute the Pho92-Not1 complex using recombinant proteins. However, for all the Pho92 domains tested, we observed a strong tendency for the complex to dissociate during size-exclusion chromatography, consistent with the moderate affinity measured between the proteins (**Fig. 3C**). In parallel, we were unable to obtain crystals of the complex between Not1(1343–1565) domain and the non-fluorescent Pho92(21–44) peptide. To stabilize the interaction between Not1 and Pho92 domains, we generated two chimeric proteins by fusing the Pho92(21–44) peptide either at the N- or C-terminus of Not1(1343–1565) domain, separated by a flexible (GSS)_5_ linker (**Fig. 4A**), a strategy successfully applied to other low affinity complexes (52). Fluorescence anisotropy experiments revealed that the Pho92(21–44)-(GSS)_5_-Not1(1343–1565) fusion (hereafter referred to as Pho92-(GSS)_5_-Not1) does not interact with the fluorescent Pho92(21–44) peptide. In contrast, the Not1(1343–1565)-(GSS)_5_-Pho92(21–44) fusion (hereafter referred to as Not1-(GSS)_5_-Pho92) does interact, albeit with a lower Kd than the untagged Not1(1343–1565) domain alone (15.75 µM *vs* 5.13 µM; **Fig. 4A**). This strongly suggests that the Pho92(21–44) region fused to the N-terminus of the Not1 domain in the Pho92-(GSS)_5_-Not1 construct, masks the binding site for the fluorescent Pho92(21–44) peptide on the surface of Not1(1343–1565). Thus, in this construct, Pho92 likely binds in *cis* with Not1, mimicking its *trans* interaction *in vivo*.

**Figure 4.**
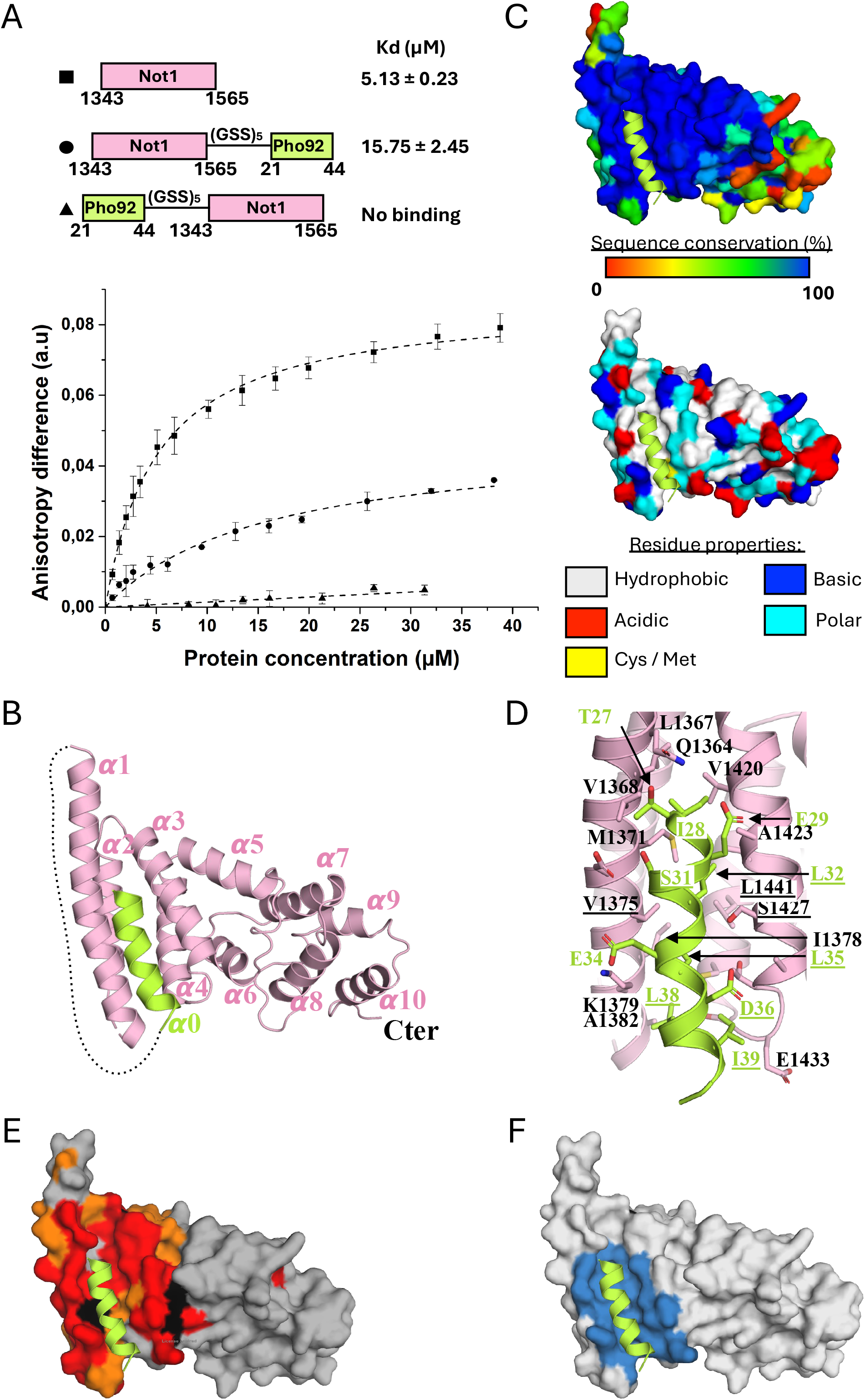
Structure of the complex between Pho92(21–44) and Not1(1343–1565) **A.** Fluorescence anisotropy analyses of the binding of Not1(1343–1565) domain or Pho92-Not1 fusion proteins and the FITC-labeled Pho92(21–44) peptide. The peptide was incubated with increasing concentrations of purified recombinant *S. cerevisiae* Not1(1343–1565) domain or Pho92-Not1 fusion proteins. The graph shows the difference in anisotropy between free fluorescent peptide and the reaction mixture as a function of protein concentration. Dashed lines represent fitted curves obtained after fitting of the experimental data as described in Materials and Methods. Error bars represent standard deviation from triplicate experiments. Kd values determined from the fits are provided alongside a schematic representation of the Not1 domains used in this experiment. **B.** Crystal structure of the Pho92(21–44)-(GSS)_5_-Not1(1343–1565) fusion construct. Pho92 and Not1 regions are colored in light green and pink, respectively. In all panels, Pho92(21–44) helix is highlighted in light green. **C.** Sequence conservation (top; calculated from an alignment of 26 *Saccharomycetaceae* proteins) and amino acid distribution (bottom) mapping at the surface of the *S. cerevisiae* Not1(1343–1565) domain. **D.** Close-up view of the interface between Pho92(21–44) and Not1(1343–1565) regions. Pho92 residues are labeled in light green and residues mutated in this study are underlined. **E.** Mapping of the Pho92(21–44) binding site on Not1(1343–1565) using NMR spectroscopy. ^1^H-^15^N HSQC NMR spectra were recorded in the absence or presence of a 1.5 molar excess of Pho92(21–44) peptide. Residue color coding: Amino acids with chemical shift peaks disappearing upon peptide addition (black); amino acids with chemical shift perturbations > 0.05 ppm and/or peak intensities < 0.35 times that of the free Not1 domain (red); amino acids with chemical shift perturbations 0.02-0.05 ppm and/or peak intensities 0.35-0.5 times that of the free Not1 domain (orange); other amino acids (grey). **F.** Surface representation of the Not1(1343–1565) domain highlighting the Pho92(21–44) fingerprint. Residues from Not1(1343–1565) directly contacting Pho92(21–44) are colored in blue.

We obtained crystals of the Pho92-(GSS)_5_-Not1 construct and determined and refined its structure at 2.68 Å resolution (**Supplementary Table S5**). Eight Pho92-(GSS)_5_-Not1 fusion proteins are present in the asymmetric unit, but only four have most of their amino acids well defined in the 2Fo-Fc electron density map. For these protomers, the regions corresponding to Pho92 residues 27-41 and Not1 residues 1352-1384 and 1392-1560 could be modeled. In contrast, for the other four protomers, the Pho92 region and Not1 residues 1343-1411 could not be modeled in the electron density. These observations suggest that the N-terminal part of this Not1 domain exhibits local flexibility (see below). In all protomers, the (GSS)_5_ linker is not visible, which is not surprising as this sequence was selected for its intrinsic high flexibility to allow the Pho92 region to interact in *cis* with the Not1 domain. Since the modeled regions from these eight Not1 protomers are virtually identical (root-mean-square deviation (RMSD) lower than 0.24 Å over at least 130 C*a* atoms) and considering that the Pho92 region binds to the same site on Not1, we will use the structure of one Pho92-Not1 complex to describe the interface.

The Not1(1343–1565) domain adopts a solenoid structure composed of five pairs of anti-parallel α-helices (α1 to α10) stacked upon each other, thus belonging to the HEAT repeat family (**Fig. 4B, Fig S2**). This domain exhibits an mIF4G fold and shares strong structural similarity with the poorly characterized central mIF4G domain from *Chaetomium thermophilum* Not1 ((53); DALI Z-score of 13, RMSD of 4.3 Å and 18% sequence identity (54). It also resembles the middle domain of human eIF4GII (Z-score of 9.4, RMSD of 4 Å and 9% sequence identity (55)) and the mIF4G domain of the human pre-mRNA splicing factor CWC22 homolog (Z-score of 9, RMSD of 4.6 Å and 16% sequence identity; (56)). A structural comparison of the *S. cerevisiae* Not1(1343–1565) domain and the central mIF4G from *Chaetomium thermophilum* Not1 reveals that their N-terminal (helices α1-α6) and C-terminal (helices α7-α10) halves are similar to each other, with RMSD values of approximately 2 Å. However, their relative orientations differ between *S. cerevisiae* and *C. thermophilum* proteins due to a 50° rotation between the two halves (**Fig. S3**).

In the complex, the Pho92 region encompassing amino acids 27-41 folds as an amphipathic α-helix (α0) that binds to a shallow groove formed by helices α1 and α2 from the Not1 domain (**Fig. 4B**). This interaction is primarily mediated by highly conserved hydrophobic residues from both proteins and has a buried surface area of 630 Å^2^, comparable to that of most protein-peptide complexes (**Fig. 4C-D**; (57)). Three leucine residues (L32, L35, L38) and two isoleucine (I29 and I39) from the hydrophobic face of Pho92 α-helix interact with numerous hydrophobic residues in Not, includingL1367, V1368, M1371, V1375, I1378, V1420, V1424, L1436, and L1441 (**Fig. 4D**, and **Fig. S2**).

In summary, our crystal structure of the Pho92-(GSS)_5_-Not1 fusion protein reveals that the Pho92(21–44) region interacts with a strongly conserved and hydrophobic groove at the surface of the Not1(1343–1565) domain. To our knowledge, this makes Pho92 the first identified binding partner of this Not1 domain.

### Validating the Pho92(21–44)-Not1(1343–1565) interface

Since a crystal structure could only be obtained by fusing Pho92 peptide to Not1, we performed chemical shift perturbation mapping using NMR spectroscopy measurements to validate that the Pho92-Not1 interface observed is not artifactual. First, we recorded ^1^H-^15^N HSQC, as well as HNCO, HNCA, HN(CO)CA, HN(CO)CACB, HN(CA)CO and HNCACB spectra for the ^13^C-^15^N-labelled Not1(1343–1565) domain alone and assigned peaks corresponding to the backbone atoms for nearly 90% of the amino acids (**Fig. S4**). The unassigned regions are primarily located in the C-terminal half of the Not1(1343–1565) domain (*i.e.* downstream of residue Ser1469) and are thus positioned opposite to the region interacting with Pho92 in our crystal structure. We noticed that sharp peaks appeared over time in the ^1^H-^15^N HSQC spectrum, indicating that the Not1(1343–1565) domain undergoes some degradation. Additionally, we observed the appearance of duplicated peaks with similar amplitudes, likely due to either proline isomerization or distinct conformations of certain Not1 regions (**Fig. S5A**). These latter peaks predominantly correspond to amino acids in the N-terminal part of this domain (**Fig. S5B**), which may explain why helices α1 and α2 from four protomers are not visible in our crystal structure.

Next, we added a 1.5-fold molar excess of the Pho92(21–44) peptide and observed chemical shift variation or peak disappearances, primarily for residues in the N-terminal part of the Not1(1343–1565) domain (**Fig. 4E, Fig. S6-8**). The Not1 residues, whose chemical environment changes in the presence of the peptide, form a large region that matches the interface observed in our Pho92-(GSS)_5_-Not1 crystal structure (**Fig. 4F**). However, as often observed in such experiments, the region affected in the NMR spectra is significantly larger than the Not1 region directly interacting with the Pho92(21–44) peptide, likely due to long-range effects (**Fig. 4E-F**).

To further validate this interface, we mutated several hydrophobic residues directly involved in the Not1-Pho92 interaction. We generated and purified the Pho92(1–69) L32R, L35R, L32R/L35R and L38R/I39R variants. All these mutants exhibited identical purification profile, indicating that these mutations do not affect the overall folding of this Pho92 construct. Using isothermal titration calorimetry (ITC), we tested their ability to interact with the Not1(1343–1565) domain. While the wild-type Pho92(1–69) domain interacts with Not1(1343–1565) domain with a Kd of 26.6 µM, the Pho92(1–69) L32R or L35R single mutants, as well as the L32R/L35R double mutant, showed no detectable interaction with Not1(1343–1565) domain under our experimental conditions (**Table 1, Fig. S9A**). The L38R/I39R double mutant still interacted but with a significantly reduced affinity Kd of 81.7 µM (**Table 1, Fig. S9A**). In parallel, we investigated whether the wild-type Pho92(1–69) or the L32R/L35R double mutant could displace the wild-type Pho92(21–44) fluorescent peptide bound to the Not1(1343–1565) domain. As expected from our ITC data and crystal structure, the wild-type Pho92(1–69) protein, but not the L32R/L35R double mutant, competed with the wild-type Pho92(21–44) fluorescent peptide for binding to this Not1 domain (**Fig. S9B**). The wild-type Pho92(1–69) peptide exhibited a Kd of 8.78 µM, compared to 4.42 µM for the Pho92(21–44) fluorescent peptide.

**Table 1:** Thermodynamics parameters for Pho92 (1–69) (WT and mutants) binding to Not1(1343–1565) at 20°C as determined by ITC.

|  | **K<sub>d</sub>**<br>**(μM)** | **Stoichiometry**<br>**(n)** | **ΔH**<br>**(kcal/mol)** | **TΔS**<br>**(kcal/mol)** |
| --- | --- | --- | --- | --- |
| **Pho92(1-69) WT<sup>a</sup>** | 26.6 ± 4.8 | 0.99 ± 0.04 | -7.51 ± 0.69 | 1.37 ± 0.59 |
| **Pho92(1-69) L32R** | ND | ND | ND | ND |
| **Pho92(1-69) L35R** | ND | ND | ND | ND |
| **Pho92(1-69)**<br>**L32R/L35R** | ND | ND | ND | ND |
| **Pho92(1-69)**<br>**L38R/I39R<sup>a</sup>** | 81.6 ± 15.4 | 0.92 ± 0.01 | -9.12 ± 1.11 | 3.63 ± 1.23 |

ND: Not Detected in two independent experiments.
<sup>a</sup> Mean values were calculated from three (WT) or two (L38R/I39R) independent experiments and standard error of the mean (SEM) are indicated.

In parallel, we also purified the Not1(1343–1565) V1375R and S1427R/L1441R mutant proteins, as the substituted residues are located at the interface with Pho92 in our crystal structure. The purification and thermostability profiles of these two mutants were broadly similar to those of the wild-type Not1(1343–1565) domain, supporting their correct folding. Again, our ITC experiments revealed that neither mutant was able to interact with Pho92(1–69) (**Table 2, Fig. S9A**).

**Table 2:** Thermodynamics parameters for Not1 (1343–1565) (WT and mutants) to binding Pho92(1–69) at 20°C as determined by ITC.

|  | **K<sub>d</sub> (μM)** | **Stoichiometry (n)** | **ΔH (kcal/mol)** | **TΔS (kcal/mol)** |
| --- | --- | --- | --- | --- |
| **Not1(1343-1565)**<br>**WT<sup>a</sup>** | 26.6 ± 4.8 | 0.99 ± 0.04 | -7.51 ± 0.69 | 1.37 ± 0.59 |
| **Not1(1343-1565)**<br>**V1375R<sup>b</sup>** | ND | ND | ND | ND |
| **Not1(1343-1565)**<br>**S1427R/L1441R<sup>b</sup>** | ND | ND | ND | ND |

ND: Not Detected in two independent experiments.
<sup>a</sup> Mean values were calculated from three independent experiments and standard error of the mean (SEM) are indicated.
<sup>b</sup> Two independent experiments indicated the absence of interaction.

Altogether, the combination of NMR data and mutagenesis experiments confirms that the Pho92-Not1 interface observed in our crystal structure of the Pho92(21–44)-(GSS)_5_-Not1(1343–1565) fusion protein is valid.

### Pho92(21–44) also interacts with the CC domain of Not1

Our fluorescence anisotropy experiments revealed that the Not1(1343–1565) domain is not the only region that directly interacts with Pho92(21–44) peptide. Indeed, the adjacent upstream domain (Not1(1071–1282)), which binds the Caf40 subunit, also interacts with it (**Fig. 3C**). To obtain structural insights on the binding mode between Pho92 and this Not1 domain, we employed the same strategies used for the Pho92/Not1(1343–1565) complex, including complex reconstitution and fusion of the two protein domains. However, we were unable to obtain crystals. We then took advantage of the recent advances in protein-protein complexes structure prediction by AlphaFold 3 (AF3; (58)), to generate a model of the Pho92(1–69) region bound to Not1(1071–1282) (**Fig. 5A**). Interestingly, the five generated models were highly consistent. In all of them, the Pho92 region encompassing residues 14 to 52 folds as two parallel helices that lie onto the CC domain, on the face opposite to the region involved in Caf40 binding (51). In these models, the L32 and L35 side chains from Pho92 are facing hydrophobic residues from Not1 (V1100, L1104, L1105 and L1200) as well as the aliphatic chain of R1101 (**Fig. 5A**). Based on these observations, we performed fluorescence anisotropy measurements to investigate whether the L32R/L35R double mutant could displace the Pho92(21–44) fluorescent peptide bound to Not1(1071–1282) domain. Similarly to our observations for the Not1(1343–1565) domain, the Pho92(1–69) L32R/L35R double mutant failed to displace the Pho92(21–44) fluorescent peptide, whereas the wild-type protein did (Kd of 14.78 µM *vs* 25.1 µM for the Pho92(21–44) fluorescent peptide; **Fig. 5B**). These results agree with the AF3 model and support the notion that the same face of the Pho92 amphipathic helix is capable of interacting with two contiguous regions from Not1.

**Figure 5.**
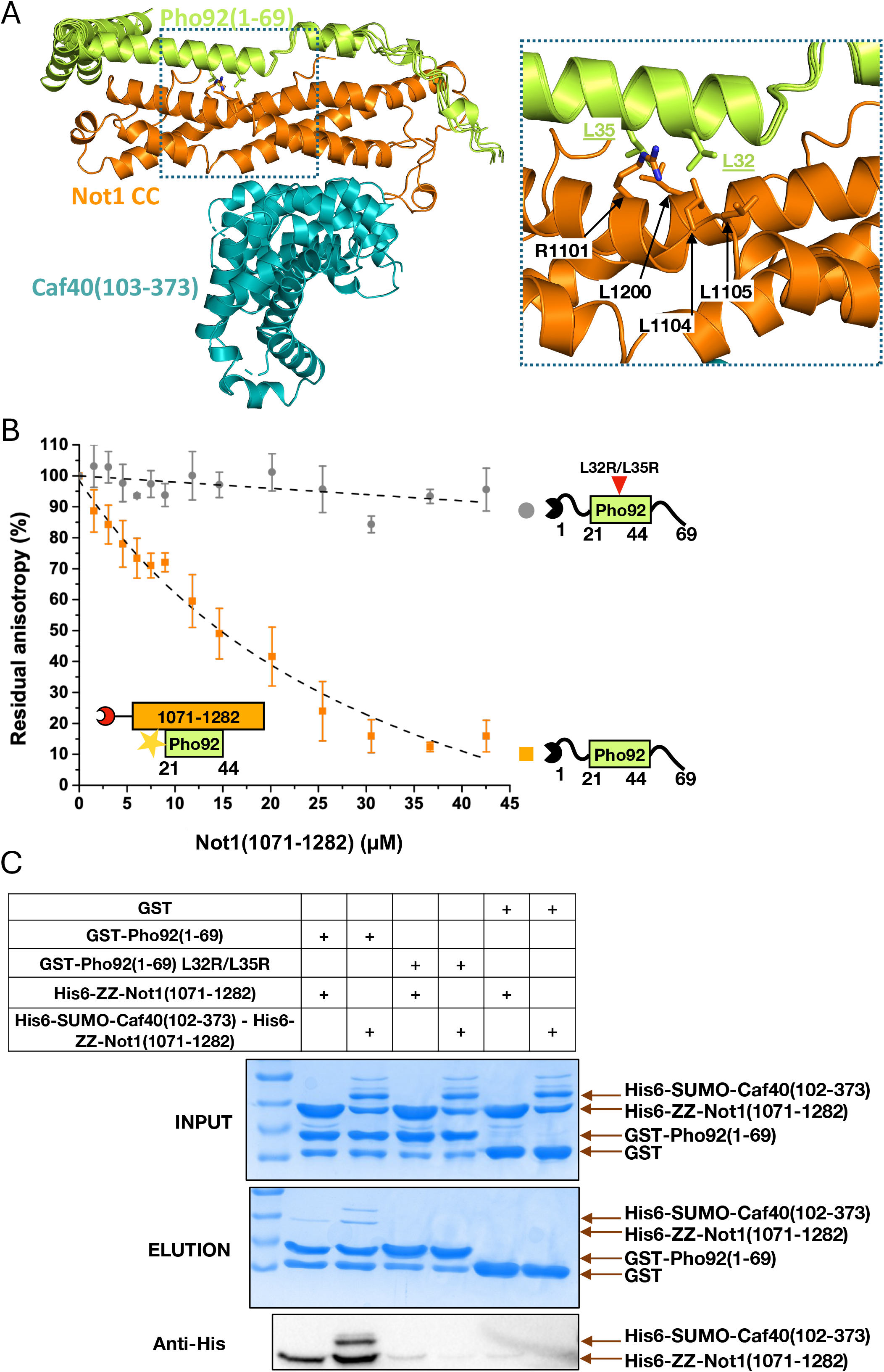
The same face of Pho92(21–44) helix interacts with Not1 CC domain. **A.** Superposition of the five AF3 models of the Pho92(1–69)/Not1(1071–1282) complex. For clarity, only one Not1(1071–1282) model is shown in orange, while the five Pho92(1–69) models are depicted in light green. The Caf40(102–373) domain (blue), bound to Not1(1071–1282), was modeled by superimposing the crystal structure of the Caf40(102–373)/Not1(1071–1282) complex (PDB code: 4CV5). Close-up view of the predicted interface with the L32 and L35 side chains from Pho92(1–69) and few Not1 residues predicted to be involved in Pho92 binding shown as sticks. **B.** Fluorescence anisotropy chase experiments assessing the displacement of FITC-labeled Pho92(21–44) bound to Not1(1071–1282) by GST-Pho92(1–69) WT (orange) or the L32R/L35R double mutant (grey). The fitted curve (dashed black line) was obtained using equation (B) from the Materials and Methods section. Error bars represent standard deviation from triplicate experiments. **C.** GST pull-down assay showing that Pho92(1–69) binds to the Not1(1071–1282)-Caf40(102–373) complex. Input and elution samples were analyzed on a 15% SDS-PAGE colored by Coomassie Blue. The presence of Not1(1071–1282) and Caf40(102–373) in the elution fractions was checked by western blot using anti-His antibodies. Three biologically independent experiments were performed, and a representative image is shown.

Since the AF3 model suggested that Caf40 and Pho92 could simultaneously interact with Not1 CC domain (**Fig. 5A**), we have co-purified the Caf40(102–373) domain together with the Not1 CC domain and tested whether this complex could interact with GST-Pho92(1–69) using GST pull-down. As controls, we confirmed that Not1 CC domain specifically interacts with GST-Pho92(1–69) but not with GST or GST-Pho92(1–69) L32R/L35R (**Fig. 5C**), consistent with our fluorescence experiments (**Fig. 3C and 5B**). Interestingly, when the Caf40(102–373)-Not1 CC complex was incubated with GST-Pho92(1–69) WT, both proteins were retained, indicating that the binding of Caf40(102–373) to Not1 CC does not prevent the interaction of this latter with Pho92, as suggested by our model (**Fig. 5A**). Furthermore, the Caf40(102–373)-Not1 CC complex did not interact with GST-Pho92(1–69) L32R/L35R, indicating that Caf40(102–373) is not directly engaged in a binding with this region of Pho92.

Thus, in addition to the Not1(1343–1565) domain, Pho92 also interacts with the Not1 CC domain without disrupting its interaction with Caf40, suggesting that Pho92 can recruit a functional CCR4-NOT complex.

### Mutations of the Pho92 conserved N-terminal motif block interaction with Not1 in vivo

To test whether mutations of the amphipathic α-helix affect the Pho92-Not1 interaction *in vivo*, we introduced the S31A-L32K and the L35K-D36A substitutions in the Pho92(1–46) construct and assessed if they abrogated its interaction with Not1 in the two-hybrid assay. We observed that both mutant proteins reduced the signal to background levels when tested against full-length Not1, as well as with the Not1(1080–2108) and Not1(1071–1565) fragments (**Fig. 6A**).

**Figure 6.**
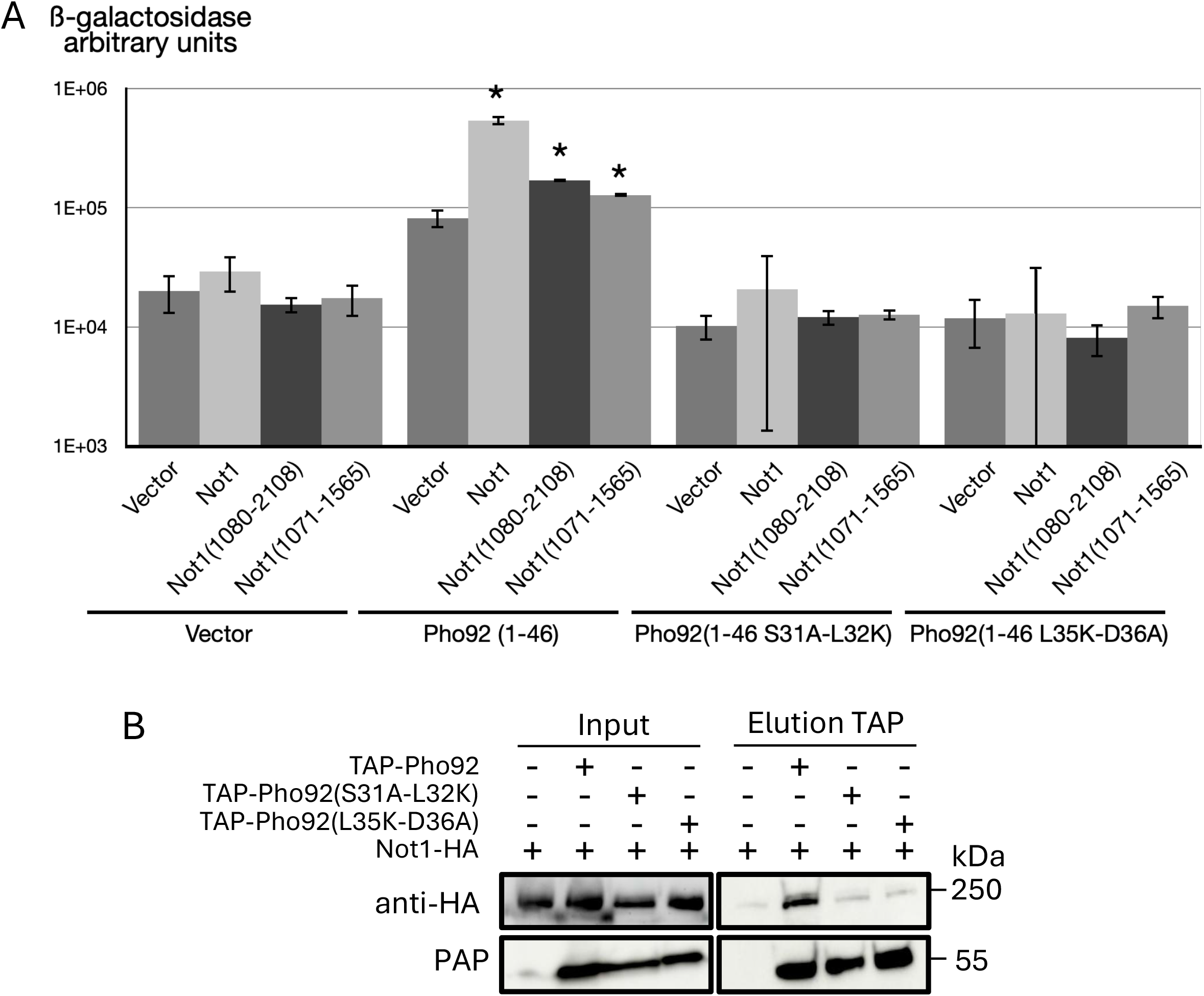
Validation of the interaction site between Pho92 motif and Not1 *in vivo*. **A.** Interaction between Pho92 (WT and mutants) and Not1 domains monitored using a yeast two-hybrid assay. Pho92 and Not1 were fused to the DBD and AD domains respectively. The interaction was quantified by measuring the β-galactosidase activity. Assays were performed as described in Figure 3A. **B.** Co-immunoprecipitation assay showing interaction between Not1 and Pho92 proteins. Protein extracts (input) were immunoprecipitated with IgG Dynabeads and resolved by SDS-PAGE. Not1-HA and TAP-Pho92 were detected using anti-HA and PAP antibodies, respectively. A representative result from multiple independent experiments is shown.

To determine whether these mutations also prevent Pho92-Not1 interaction during meiosis, we constructed strains expressing the mutated TAP-tagged Pho92 from its chromosomal locus using CRISPR-Cas9 in a background expressing HA-tagged Not1. We then prepared protein extracts from these cells at 5 hours of meiosis and immunoprecipitated TAP-Pho92 using IgG beads. Analysis of the immunoprecipitated proteins confirmed that Not1 co-immunoprecipitated by wild-type Pho92 and demonstrated that mutations of the interface residues of the Pho92 N-terminal amphipathic α-helix abolish this association (**Fig. 6B**). Overall, these results are consistent with our biochemical assays (**Fig. 3 and 5**) demonstrating that the N-terminus of Pho92 mediates its interaction with Not1 *in viv*o. Moreover, even if multiple Pho92 binding sites coexist on Not1, these substitutions abolish all interactions.

## Discussion

Our analyses demonstrate that a conserved present near the N-terminus of *S. cerevisiae* Pho92 binds Not1 and recruits the CCR4-NOT deadenylase. This is consistent with our observation of m^6^A-dependent deadenylation during meiosis. The conserved residues fold into an amphipathic helix that directly interacts with two subdomains of Not1. At the first glance, the observation that a single motif binds to different Not1 regions may seem surprising. However, this is a recurrent theme among proteins that recruit the main deadenylase, as several of them have been shown to interact with different sites on Not1 (2). The predominantly α-helical structure of Not1, possibly resulting from internal duplication during evolution, may explain some of these observations. Interestingly, the Not1 domain with the highest affinity for Pho92 had not previously been shown to interact with any partner. This domain adopts an mIF4G fold, many of which have been previously shown implicated in protein-protein interactions, particularly with RNA helicases. Given that amphipathic helices equivalent to the Pho92(21–44) region are found in other proteins, it is possible that additional factors use a similar mode of interaction to associate with this domain of Not1.

Our observation that two Not1 domains can interact with the same region of Pho92 suggests that the presence of multiple m^6^A sites on an mRNA, as observed on some yeast mRNAs (27, 28), could synergistically enhance CCR4-NOT complex recruitment, thereby potentiating rapid deadenylation and decay (**Fig. 7**). However, contrary to what was previously reported for human mRNAs (17), to our knowledge, there is no evidence for a correlation between the number of m^6^A sites on yeast mRNAs and their kinetic of degradation.

**Figure 7.**
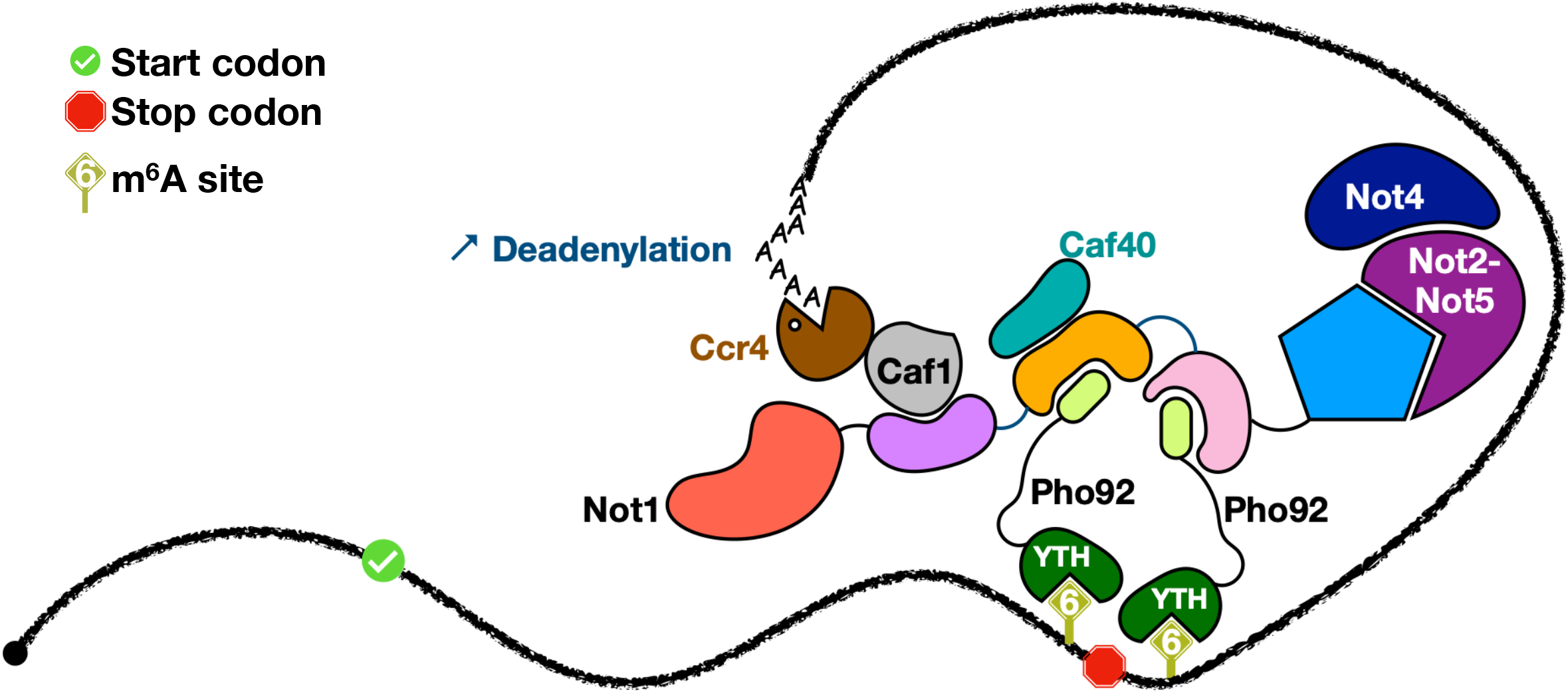
Model of the CCR4-NOT complex recruitment by multiple Pho92. Not1 is schematized with its different globular domains, linked by a line. Associated CCR4-NOT subunits and two Pho92 proteins are also depicted, the latter bound to two m6A residues present in the same mRNA, resulting in possible synergistic amplification of the interactions to stimulate deadenylation.

Altogether, our observations further expand the number of Not1 sites involved in partner interactions, reinforcing the concept that the CCR4-NOT scaffold protein Not1 acts as a giant adaptor to connect this complex with proteins that target it to specific substrates (2). Previous co-immunoprecipitation studies had suggested that Pho92 physically interacts with the Caf1 deadenylase subunit of the CCR4-NOT complex (29). However, these observations were questionable as they were performed under conditions in which Pho92 is not naturally expressed. Our data indicate that the interaction between Caf1 and Pho92 is indirect and mediated by Not1 (**Fig. 3A**).

Interestingly, the motif present in Pho92 is conserved in some *Saccharomycetaceae* species, but no related region is found in the human YTHDF homologs. This may reflect the the rapid evolution of such short linear motifs compared to the more constrained evolution of globular domains. Notably, this feature may enable a rapid rewiring of protein interaction networks (59), as previously observed for the yeast specific interaction of the mRNA decay factors Xrn1 and Dcp2 with Pat1 decapping activator (60). In fact, although the Pho92 motif is conserved in some *Saccharomycetaceae* species, careful analyses indicate that it is absent from certain members of this family, including the *Kluyveromyces* phylum. Like human YTHDC proteins, *Kluyveromyces* Pho92 contains conserved motifs of different identity N-terminal to its YTH-domain. Whether these motifs are involved in the interaction with the CCR4-NOT complex or other cellular factors remains unknown. In the case of the human YTHDF2 protein, experimental evidences have suggested a role for the unstructured N-terminal region of YTHDF2 in interacting with the CCR4-NOT complex (14, 61). One of these studies concluded that residues 101-200 interacted directly with the CCR4-NOT complex (14). However, this conclusion was based on experiments lacking appropriate controls: the data presented did not exclude the possibility that YTHDF2(101–200) nonspecifically bound to the resin during the pull-down experiment. Furthermore, the levels of recovered YTHDF2(101–200) were in high molar excess over Not1 fragment used for the pull-down (14, 62, 63), further questioning whether a biologically relevant interaction was detected. Thus, while YTHDF2 was described in the literature as associating with the CCR4-NOT complex, further analyses will be required to determine whether it binds CNOT1 directly. If this is the case, it would be of interest to identify the specific residues of YTHDF2 and CNOT1 involved in this interaction.

Interestingly, the *Schizosaccharomyces pombe* YTH domain containing protein Mmi1 has also been shown to interact with the CCR4-NOT complex (62, 63). Although it has lost its ability to recognize m^6^A marks (64), consistent with this organism lacking m^6^A modifications, its YTH domain has acquired the capacity to recognize specific RNA sequences through a novel interaction surface. Like its *S. cerevisiae* Pho92 counterpart, Mmi1 is also involved in the regulation of meiotic mRNAs. However, while Pho92 functions during meiosis, Mmi1 eliminates meiotic mRNAs during vegetative growth. Although Mmi1 lacks the conserved motif present in Pho92, it still maintains its association with the CCR4-NOT complex through an as yet uncharacterized interface (62). Further, the region of Mmi1 N-terminal to the YTH domain has acquired the ability to interact with other factors, including Erh1, which is also involved in the downregulation of meiotic transcripts during vegetative growth (52, 65). Once again, this illustrates the rapid rewiring of protein interaction networks, particularly in cases involving YTH proteins.

Our data demonstrate that the Pho92 interaction with the CCR4-NOT complex is critical for meiosis. Previous studies had suggested that Pho92 recruits the CCR4-NOT complex to m^6^A-containing mRNAs, but molecular mechanistic details were lacking (28). Our results provide a firm structural basis for the tethering of the CCR4-NOT complex to m6A containing mRNAs. Since previous transcriptomic analyses have shown that m^6^A-containing mRNAs persist in the absence of Pho92 (27), it is likely that their deadenylation initiates their degradation.

Altogether, our observations reveal how Pho92 interacts with the main deadenylase CCR4-NOT to promote meiotic completion in the yeast *Saccharomyces cerevisiae*. Further analyses will be required to dissect whether this interaction is constitutive or whether it is finely tuned to occur exclusively at specific stage of the yeast meiotic program.

## Acknowledgements

We thank members of our teams for critical reading of the manuscript and suggestions. We are also grateful to Ed Hurt for the HO plasmid, M. Rougemaille for technical suggestions, F. Wyers for the anti-Stm1 antibody, and F. van Werven for kindly sharing the pCUP-IME1 strain, and IGBMC platforms particularly the Imaging, Mass Spectrometry, Peptide Synthesis and Mediaprep facilities for support. We acknowledge the French TGIR-RMN for supporting this project by giving free access to its infrastructure. We acknowledge SOLEIL for provision of synchrotron radiation facilities and particularly staff from Proxima-2A beamline.

## Conflict of interest

There are no conflicts to declare.

## Funding

This work was supported by the Agence Nationale de la Recherche grant ANR-16-CE11-0003 [to M.G. and B.S.], ANR-21-CE12-0029 [to M.G.] and ANR-22-CE11-0015 [to M.G. and B.S.], the CNRS [to M.G.], Ecole Polytechnique [to M.G.] and the IGBMC [to B.S.]. This work of the Interdisciplinary Thematic Institute IMCBio, as part of the ITI 2021-2028 program of the University of Strasbourg, CNRS and Inserm, was supported by IdEx Unistra (ANR-10-IDEX-0002), by SFRI-STRAT’US project (ANR 20-SFRI-0012), and EUR IMCBio (ANR-17-EURE-0023) under the framework of the Plan France 2030. D. H. was supported by a PhD fellowship from the French Ministère de l’Enseignement Superieur et de la Recherche (MESR).

## Author contributions

Experimental planning, writing of original draft, figure preparation and manuscript reviewing, L.S., D.H., J.S., N.M., B.S. and M.G.; experiment realization L.S., D.H., J.S., N.M., N.U., M.M., H.B., M.R., S.A., and M.G.; project supervision, C.vH., B.S. and M.G.; funding, M.G. and B.S.

## Material availability

Described material is available from the corresponding authors upon reasonable request.

## Notes

### Competing Interest Statement

The authors have declared no competing interest.

